# Mesenchyme governs hair follicle induction

**DOI:** 10.1101/2023.06.29.547048

**Authors:** Otto J.M. Mäkelä, Marja L. Mikkola

**Author notes:** Corresponding author: Marja L. Mikkola.

## Abstract

Tissue interactions are essential to guide organogenesis. The development of hair follicles is regulated by inductive signalling between the embryonic surface epithelium and the adjacent mesenchyme. Previous studies have established that the mesenchymal component of the hair follicle, the dermal papilla and its precursor dermal condensate, has the capacity for de novo hair follicle induction. However, studies prior to dermal condensate formation have been inconclusive, and therefore the source and the identity of the primary inductive signal have remained unknown. Here, we performed epithelial-mesenchymal tissue recombination experiments using hair-forming back skin and glabrous plantar skin from mouse embryos to unveil that the back skin mesenchyme is inductive even prior to dermal condensate formation. Moreover, the naïve, unpatterned mesenchyme was sufficient to trigger hair follicle formation even in the oral epithelium. Considering the recognized role of Wnt signalling and Bmp activity inhibition in initiation of hair follicle development, we explored the hair-inductive ability of the Wnt agonist R-spondin-1 and a Bmp receptor inhibitor in embryonic skin explants. Although R-spondin-1 instigated precocious placode-specific transcriptional responses, it alone or in combination with the Bmp receptor inhibitor was insufficient for hair follicle induction. Our findings pave the way for identifying the hair follicle-inducing cue.

## Introduction

Inductive tissue interactions constitute a fundamental mechanism regulating embryonic development. As most organs develop from multiple tissue types, cell-cell communication across tissue boundaries is critical to ensure successful organ induction, patterning, morphogenesis, and cell differentiation (Thesleff, Vaahtokari & Partanen, 1995). The hair follicle is an excellent model to study these interactions as it develops from the embryonic surface ectoderm and the underlying mesenchyme, and reciprocal signalling between these compartments directs its development, as well as the life-long cyclic function of the mature organ (Hardy, 1992; Sennett & Rendl, 2012; Morgan, 2014; Li & Tumbar, 2021). The inductive signalling regulating hair follicle development and renewal has been under meticulous investigation for decades, yet some key aspects remain elusive to date.

In mice, hair follicles are induced in three consecutive waves, the first one commencing around embryonic day E13.5 (Fliniaux et al., 2008; Biggs et al., 2018) as a response to a yet unidentified primary inductive signal. As a result, epithelial keratinocytes migrate centripetally to form a thickening, the placode (Ahtiainen et al., 2014). Inductive signals from the placode then induce the underlying fibroblasts to migrate and aggregate underneath the placode, forming the mesenchymal component of the hair follicle, the dermal condensate (DC) (Glover et al., 2017; Biggs et al., 2018; Gupta et al., 2019; Mok et al., 2019). Subsequently, secondary inductive signals from the DC promote invagination of the epithelium to the mesenchyme, and the DC is engulfed by the epithelium and remains an integral part of the mature hair follicle, named the dermal papilla (DP) (Sennett & Rendl, 2012). As the embryo grows larger, secondary and tertiary hair placodes form between the existing hair primordia in two waves at around E16.5 and E18.5, respectively (Duverger & Morasso, 2009).

Tissue crosstalk between the follicular epithelium and the underlying mesenchyme is mediated by conserved signalling pathways (Biggs & Mikkola, 2014). Initially, Wnt/β-catenin signalling is focally activated in the prospective placode where it is necessary for placode formation as suppression of epithelial β-catenin signalling activity blocks hair placode formation altogether (Andl et al., 2002; Zhang et al., 2009; Huelsken et al., 2001). Moreover, forced activation of epithelial Wnt/β-catenin signalling is thus far the only known genetic manipulation to induce precocious formation of hair placodes (Närhi et al., 2008; Zhang et al., 2008). After initial epithelial Wnt activation, Eda/Edar/NF-κB pathway is necessary to reinforce Wnt activity and placode morphogenesis to continue (Schmidt-Ullrich et al., 2006; Fliniaux et al., 2008; Zhang et al., 2009). Once placode formation has initiated, signals emanating from the placode, in particular fibroblast growth factor 20 (Fgf20) and Sonic hedgehog (Shh), govern DC formation (Huh et al., 2013; Woo, Zhen & Oro, 2012). Fgf20 has been shown to induce directional migration of mesenchymal fibroblasts and to induce DC cell fate (Mok et al., 2019; Biggs et al., 2018). Fgf20 is necessary for DC formation, as demonstrated by the absence of all molecular and morphological indications of primary and secondary DCs in *Fgf20* null mice (Huh et al., 2013; Biggs et al., 2018; Mok et al., 2019).

Several lines of evidence indicate that suppression of Bmp signalling is essential for hair follicle induction. Deletion of Bmp-inhibitor gene *Noggin*, expressed in the DC, suppresses secondary and tertiary hair placode formation (Botchkarev et al., 1999, 2002), while its ectopic expression under the *Keratin 14* promoter results in increased hair follicle density and formation of ectopic hair follicles in ventral paws and at the site of the Meibomian glands (Plikus et al., 2004). Furthermore, Noggin-treatment increases hair placode density in embryonic skin explants, whereas Bmp4-treatment prevents hair placode formation by suppressing expression of *Edar* and *Lef1*, the mediator of Wnt/β-catenin pathway (Botchkarev et al., 1999; Mou et al., 2006; Jamora et al., 2003).

In the past, hair follicle induction has been studied extensively by transplantation and tissue recombination experiments. It was first demonstrated that the whisker follicle DP contains the capacity to induce new follicles by transplantation of rat vibrissae DPs in the ear skin and subsequent formation of ectopic whisker follicles (Cohen, 1961). Later, tissue recombination of hair-forming dorsal mesenchyme with the glabrous plantar epithelium revealed that the inductive capacity resides in the dorsal skin mesenchyme as early as the DC forms (Kollar, 1970). These and other studies have established that the hair follicle dermal component, the DP and its precursor the DC, contain the capacity to induce *de novo* hair follicle formation in the skin, as well as other epithelial tissue of ectodermal origin, such as dental and corneal epithelia (Oliver, 1970; Kollar, 1970; Hardy, 1992; Ferraris et al., 2000; Dhouailly, 1973; Cohen, 1961; Kollar & Baird, 1970; Higgins et al., 2013; Jahoda et al., 1992). The extrapolation of these findings has established the current paradigm, which posits that the mesenchyme is the site of the hair-inductive signal. However, experiments investigating the inductive capacity of the mesenchyme prior to DC formation are scarce. In fact, recombination of mouse back skin mesenchyme prior to hair follicle induction (E12.5) with glabrous epithelia of chick embryonic skin did not result in appendage induction (Dhouailly, 1973). In contrast, the reverse recombination of the E12.5 hair-forming back skin epithelium with the glabrous chick skin mesenchyme produced short epithelial nodules protruding into the dermis (Dhouailly, 1973). Furthermore, studies showing that DC formation is preceded by a placodal prepattern (Huh et al., 2013; Glover et al., 2017; Biggs et al., 2018; Mok et al., 2019) and that primary and secondary hair placodes form in the absence of DCs in *Fgf20* null embryos (Huh et al., 2013) have challenged the prevailing model of mesenchymal induction of hair follicle formation.

Here, we sought to identify the source of the hair follicle inductive cue by performing tissue recombination experiments between heterologous epithelia and mesenchymes using genetically labelled tissues expressing fluorescent reporters allowing unambiguous assessment of tissue purity. Our results reveal that the hair follicle inductive signals reside in the mesenchyme even prior to DC formation. We show that mesenchymal expression of the Wnt pathway agonist *R-spondin-1 (Rspo1*) is upregulated at the time of hair follicle induction. This prompted us to assess the impact of Rspo1 on hair follicle induction in *ex vivo* cultured skin explants. Our results indicate that Rspo1 alone, or in combination with inhibition of Bmp-signalling, is insufficient to induce precocious hair follicle formation, although the latter altered hair placode patterning and the former instigated hair placode-specific transcriptional responses even prior to hair follicle induction. We expect our results will motivate further research to identify the mesenchymal hair-inductive cues.

## Results

### Hair follicle inductive potential resides in the mesenchyme even prior to dermal condensate formation

To determine whether the hair follicle inductive signal arises from the epithelium or the mesenchyme, we performed heterotypic tissue recombination between the hair-forming back skin and the glabrous plantar skin by enzymatically separating the tissue compartments, recombining them, and culturing *ex vivo*. To monitor the tissue separation efficiency and retrospectively verify the origin of the cells, we collected epithelia from R26R^mTmG/+^ and mesenchymes from R26R^mG/+^ littermate embryos, ubiquitously expressing membrane-bound tdTomato (mT) and membrane-bound eGFP (mG), respectively. Alternatively, epithelia were obtained from R26R^floxedTomato/+^, or R26R^floxedTomato/+^;K17-GFP and mesenchymes from wild type embryos. First, we performed homotypic recombinations of E14.5 tissues that served as technical controls. Dorsal skin mesenchyme and epithelium cultured for 4 days readily formed hair follicle-like appendages, whereas homotypic recombinants of plantar tissues did not generate any appendages (Fig. 1A) validating our experimental set-up. In heterotypic recombinants, the DC-containing dorsal mesenchymal tissue induced hair follicle-like appendage formation in the glabrous plantar epithelium (Fig. 1A), as expected. No appendages were observed in the recombinants consisting of dorsal skin epithelium and plantar mesenchyme (Fig. 1A).

**Fig. 1.**
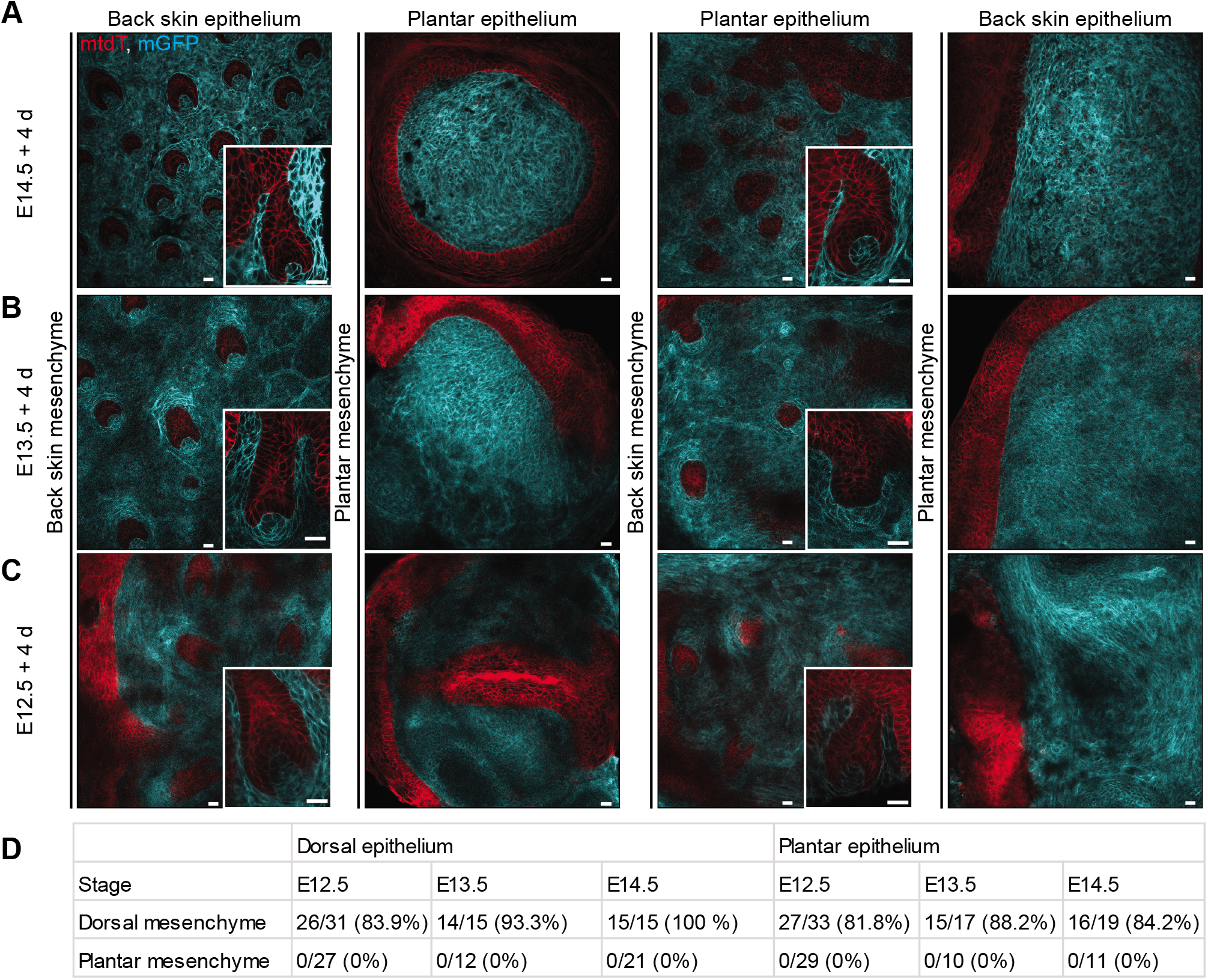
Back skin mesenchyme harbors skin appendage inductive capacity even prior to dermal condensate formation. Confocal microscopy optical sections of tissue recombinants consisting of R26R^mG/+^ (cyan) mesenchyme and R26R^mTmG/+^ (red) epithelium from embryonic back skin and plantar skin from E14.5 (A), E13.5 (B), and E12.5 (C) embryos after 4 days of *ex vivo* culture. Insets show individual appendages. Scale = 20 μm. (D) Total number of tissue recombinants of the indicated stages. Numbers indicate recombinants with appendages / total number of recombinants.

We then asked where the inductive potential resides at the time of hair placode induction at E13.5, when DCs cannot yet be morphologically discerned. Like E14.5 tissues, homotypic recombinations between back skin epithelium and mesenchyme readily produced skin appendages, whereas recombinations between plantar skin compartments did not (Fig. 1B). Interestingly, at this stage too, hair follicle-like appendages were induced upon heterotypic recombination of the dorsal mesenchyme with the plantar epithelium (Fig. 1B), while dorsal skin epithelium was not able to induce appendages in the glabrous plantar mesenchyme (Fig. 1B), indicating that the dorsal mesenchyme is inductive even prior to DC formation.

As DC formation is a gradual process and the first loosely organized Sox2+ cells are found below the emerging placodes from E13.5 onward (Biggs et al., 2018; Gupta et al., 2019), small DCs might already exist at E13.5. Therefore, we assessed where the hair follicle inductive potential resides prior to any known molecular or morphological signs of hair follicle formation by recombining tissues from E12.5 embryos. While the homotypic control recombination of the back skin formed appendages and the corresponding recombination of plantar tissues did not (Fig. 1C), we again observed appendages in the heterotypic recombinants consisting of the dorsal skin mesenchyme and glabrous plantar epithelium, but not in explants of dorsal skin epithelium and plantar mesenchyme (Fig. 1C). Quantification of all E12.5-E14.5 recombinant explants confirmed that only mesenchymal back skin was inductive (Fig 1D). These data imply that the hair-inductive potential resides in the mesenchyme, even prior to DC formation.

To characterise the appendages that formed in the E12.5 tissue recombinants, epithelia were collected from R26R^floxedTomato/+^;K17-GFP; embryos in which tdTomato (tdT) is ubiquitously expressed and the nascent hair placodes are marked by K17-GFP reporter expression (Bianchi et al., 2005) and the mesenchyme from wild type littermates. Homotypic back skin recombinations served as a positive control revealing strong K17-GFP expression in developing hair follicles (Fig. 2A). Likewise, K17-GFP expression marked the appendages formed in recombinants between back skin mesenchyme and plantar epithelium, however, recombinants of the back skin epithelium with the plantar mesenchyme were devoid of K17-GFP+ appendages (Fig. 2A). Immunostaining with the early hair follicle marker Lhx2 (Rhee, Polak & Fuchs, 2006) not expressed in sweat gland placodes (Cao et al., 2021), and the DC-marker Sox2 (Driskell et al., 2009) verified that the formed appendages were hair follicles (Fig. 2A, Fig. S1A,B).

**Fig. 2.**
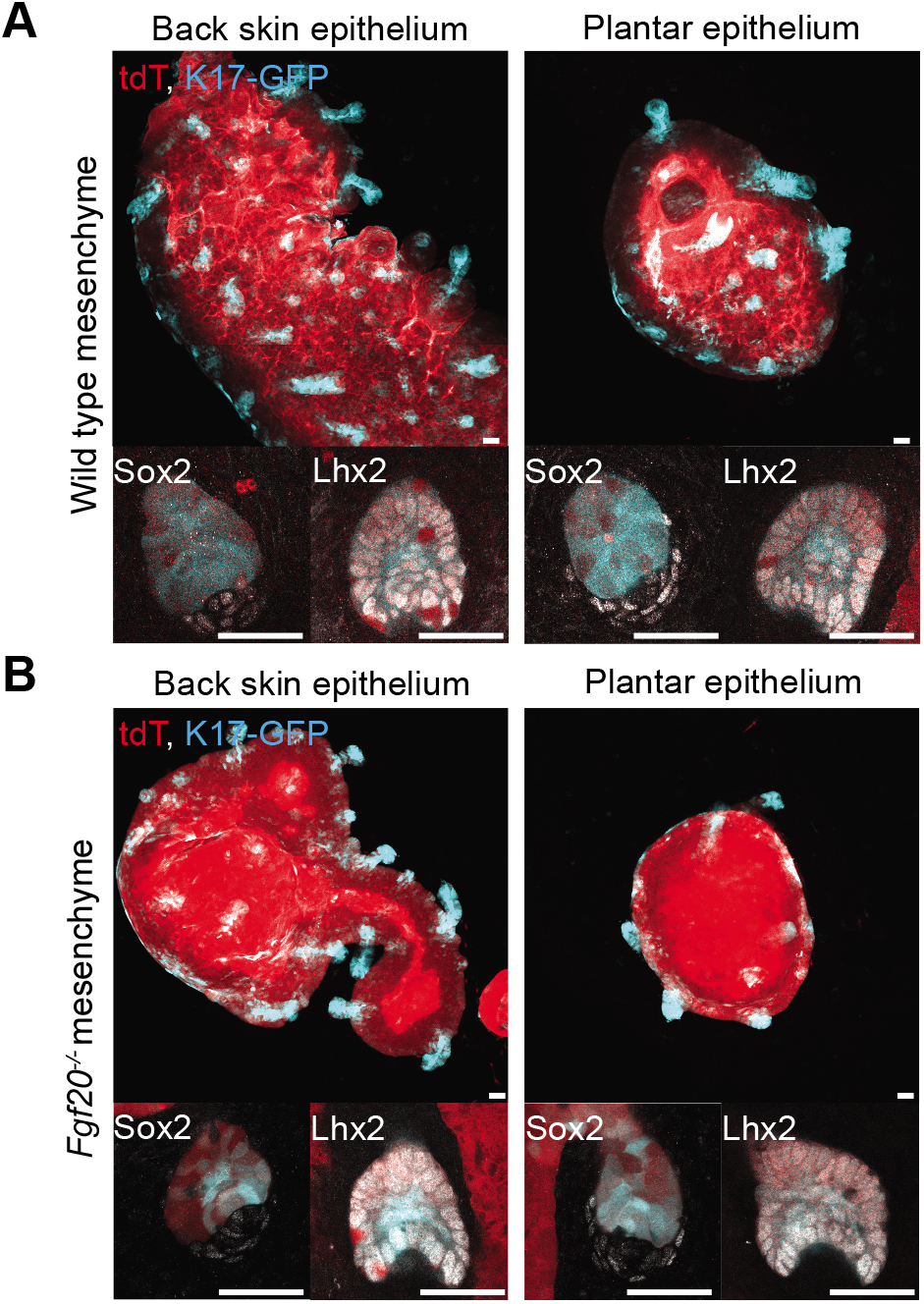
Wild type and *Fgf20* null back skin mesenchyme-induced skin appendages express hair follicle markers. (A) Maximum intensity projections of confocal microscopy images of tissue recombinants of E12.5 wild type mesenchyme and E12.5 R26R^floxedTOM^;K17-GFP epithelium from back skin (N=11/12) and plantar skin (N=11/12) (left) and optical sections of the same recombinants stained with Sox2 and Lhx2 (right). N indicates recombinants with HFs / total number of recombinants (B) Maximum intensity projections of confocal microscopy images of tissue recombinants of E12.5 Fgf20^-/-^ mesenchyme and E12.5 R26R^floxedTOM^;K17-GFP epithelium from back skin (N=15/17) and plantar skin (N=14/16) (left) and optical sections of the same recombinant stained with Sox2 and Lhx2(right). N indicates recombinants with HFs / total number of recombinants. Scale = 20 µm.

To ascertain that the mesenchyme harbours hair inductive capacity even in the absence of DC formation, we performed tissue recombination experiments using E12.5 back skin mesenchyme isolated from *Fgf20* null embryos that lack primary and secondary hair follicle DCs (Huh et al., 2013). Indeed, like the wild type mesenchyme, the *Fgf20* null mesenchyme induced formation of appendages when recombined with wild type back skin epithelium, or wild type plantar epithelium (Fig. 2B). These appendages, too, were associated with Sox2+ DCs and Lhx2+ epithelial cells (Fig. 2B, Fig. S1C,D) confirming that the hair follicle inductive capacity of the back skin mesenchyme is independent of DC formation.

Lastly, we asked whether the capacity of the naïve dorsal mesenchyme to induce appendages is restricted to epithelia of epidermal origin by assessing whether hair follicles could form in the oral epithelium. To this end, we recombined non-fluorescent E12.5 back skin mesenchyme with the epithelium of R26R^floxedTomato/+^;K17-GFP embryos, isolated from the E12.5 lower jaw diastema, a toothless region of the oral cavity between the incisor and the molar tooth primordia (Fig. S2A). Back skin epithelium from K17-GFP;R26R^floxedTomato/+^ embryos was used as a control. Again, K17-GFP positive hair follicles readily formed in homotypic back skin control recombinants after 5 days of culture (Fig. 3A). Strikingly, hair follicle-like, K17-GFP+ appendages formed also in explants consisting of the dorsal mesenchyme and the oral epithelium (Fig. 3A). Like the control explants, these appendages expressed Lhx2 and Sox2 in the epithelium and the mesenchyme, respectively. Furthermore, the recombinants generated visible hair shafts after 2 weeks of culture (Fig. 3A, Fig. S2B,C) confirming their hair follicle identity.

**Fig. 3.**
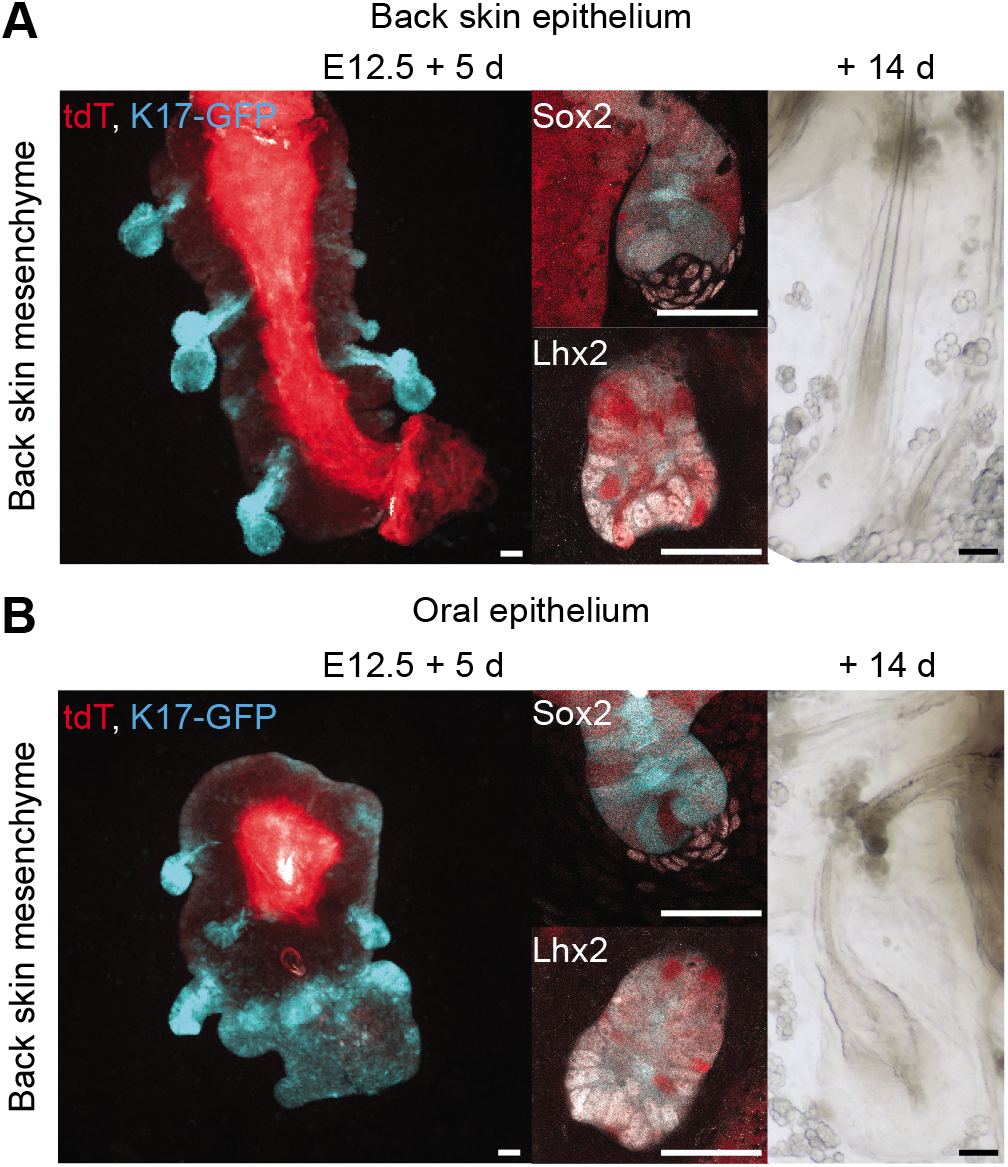
Naïve back skin mesenchyme induces hair follicles in oral epithelium. Maximum intensity projections of confocal microscopy images (left) of tissue recombinants of E12.5 wild type mesenchyme and (A) E12.5 R26R^floxedTOM^;K17-GFP back skin epithelium (N=10/12), or (B) epithelium from the toothless oral diastema region (N=13/17) after 5 days of *ex vivo* culture. N indicates recombinants with HFs / total recombinants. Optical sections of recombinants stained with Sox2 and Lhx2 (middle), and stereomicrosope images of recombinants cultured for 14 days *ex vivo* (N=5 explants) showing hair follicles producing a hair shaft (right). Scale = 20 µm.

### Mesenchymal cell density is not significantly increased prior to hair follicle induction

Next, we asked how the mesenchymal primary inductive signal is generated in the mesenchyme. As induction of feather placodes is associated with an increase of mesenchymal cell density (Wessells, 1965), we hypothesized that perhaps a similar increase of mesenchymal cell density occurs also during hair follicle induction. Given that Wnt/β-catenin signalling activity is essential for mesenchymal cells to gain DC identity (Atit et al., 2006; Gupta et al., 2019), we analysed Wnt-positive cells.

To assess mesenchymal cell density, we produced confocal 3D image stacks of transversal vibratome sections from mouse embryos prior to hair follicle induction at E12.5 and at the time of hair induction at E13.5 (Fig. 4A). We quantified the density of mesenchymal nuclei within 80 µm distance from the epithelium and calculated nuclear densities as average rolling densities in 20 µm windows at 1 µm intervals (Fig. S3A). For statistical analysis, the data was binned at 20 µm intervals. Although cell densities were significantly higher closest to the epithelium at both stages analysed, no difference was observed between E12.5 and E13.5 (Fig. 4B).

**Fig. 4.**
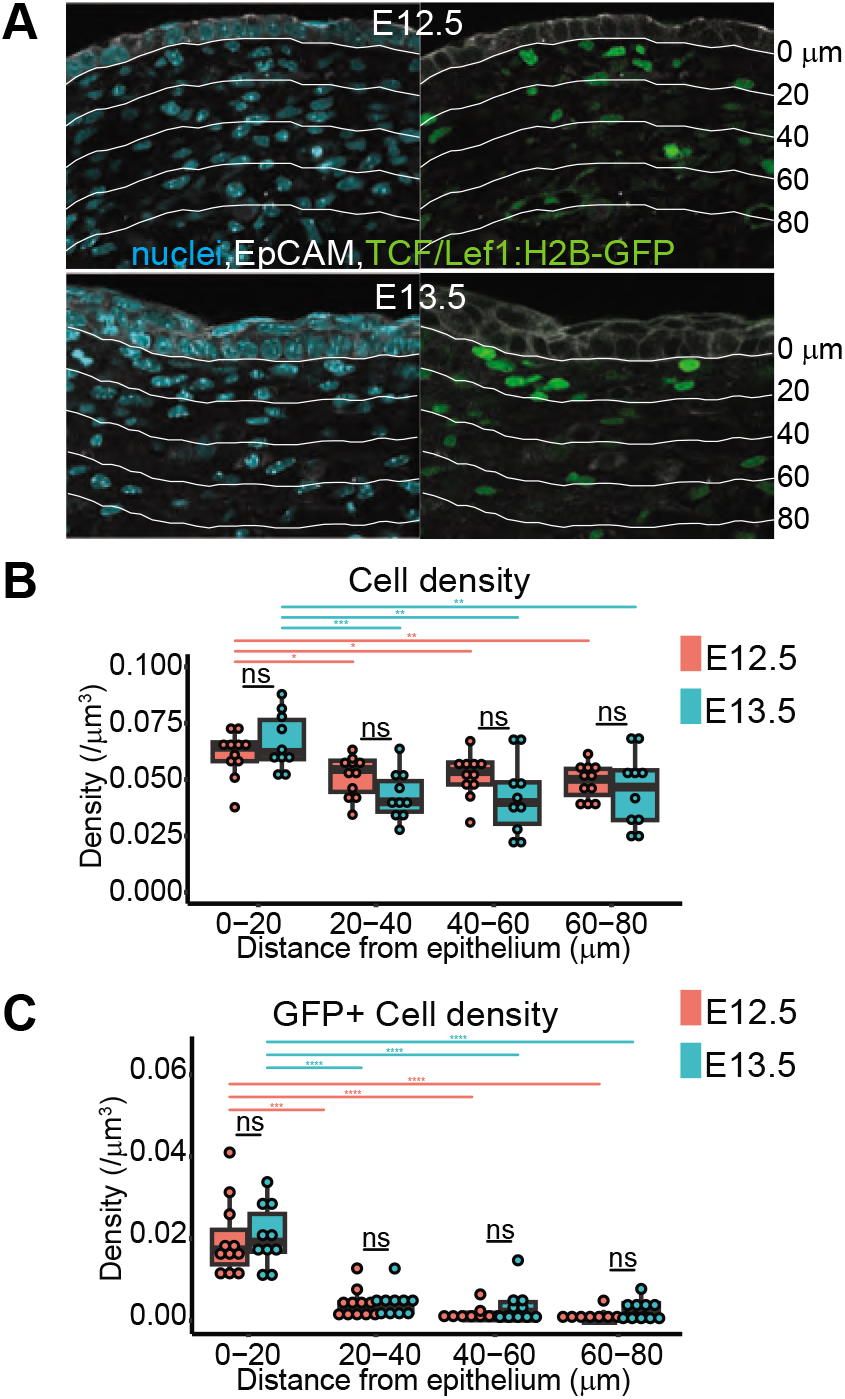
Mesenchymal cell density is unchanged during hair follicle induction. (A) Confocal optical images of transversal thoracic vibratome sections of E12.5 and E13.5 TCF/Lef1:H2B-GFP embryos stained with Hoechst33342 for nuclei and epithelial marker EpCAM. Lines indicate 20 µm distance intervals from the epithelial-mesenchymal border. Scale = 20 µm. (B) Binned cell densities in 20 µm bins (Mean ±SD) at E12.5 (red) and E13.5 (blue). (C) TCF/Lef1:H2B-GFP+ cell densities at 20 µm bins (Mean ±SD) from the epithelial-mesenhcymal border at E12.5 (red) and E13.5 (blue). N_E12.5_=11 embryos and N_E13.5_=10 embryos. Statistical significance was assessed with Student’s T-test.* P<0.05, **P<0.01, ***P<0.001.

To assess the distribution of Wnt-activated cells, we took advantage of the TCF/Lef1:H2B-GFP Wnt signalling reporter mouse (Ferrer-Vaquer et al., 2010) and calculated the average rolling density of GFP+ dermal cells at both stages (Fig. S3B). Highest density of TCF/Lef1:H2B-GFP positive cells was observed in the proximity of the epithelium, confirmed by statistical analysis of the binned data, yet there was no difference between the two stages (Fig. 4C). These results suggest that the signal initiating the formation of hair follicles likely arises through changes in gene expression of the mesenchymal fibroblasts rather than via an increase in their density or change in the proportion of Wnt/β-catenin activated cells.

### Inhibition of BMP signalling affects placode patterning, but is not sufficient to induce hair follicle development

We next aimed to gain insights on the molecular composition of the mesenchymally-derived hair follicle inductive signal using a candidate approach. The evidence showing that Bmp signalling inhibits hair placode formation (Botchkarev et al., 1999; Plikus et al., 2004; Mou et al., 2006; Pummila et al., 2007) led us to hypothesize that downregulation of BMP signalling activity could be sufficient to initiate formation of hair follicles. However, in addition to *Noggin*, numerous other Bmp antagonists are expressed in the embryonic skin mesenchyme at the time of hair follicle induction(Sennett et al., 2015; Jacob et al., 2022 preprint, http://kasperlab.org/embryonicskin) making a genetic approach challenging. Therefore, to test our hypothesis, we inhibited BMP signalling with LDN193189, a small molecule inhibitor of BMP type I receptors (Cuny et al., 2008). Back skin of E12.5 Fgf20^βGal/+^ embryos were cultured with 1 µM LDN193189 for 12 and 24h and hair placode induction was assessed by analysing the expression of hair placode markers Fgf20 (using anti-β-Gal staining as a surrogate) and Edar by whole-mount immunostaining (WMIF). Unsatisfyingly, suppression of Bmp activity did not induce precocious hair follicle formation at 12 hours (Fig. 5A). However, when placodes started to emerge after 24h culture, they were larger and less regular in size in LDN193189-treated explants compared to controls (Fig. 5B).

**Fig. 5.**
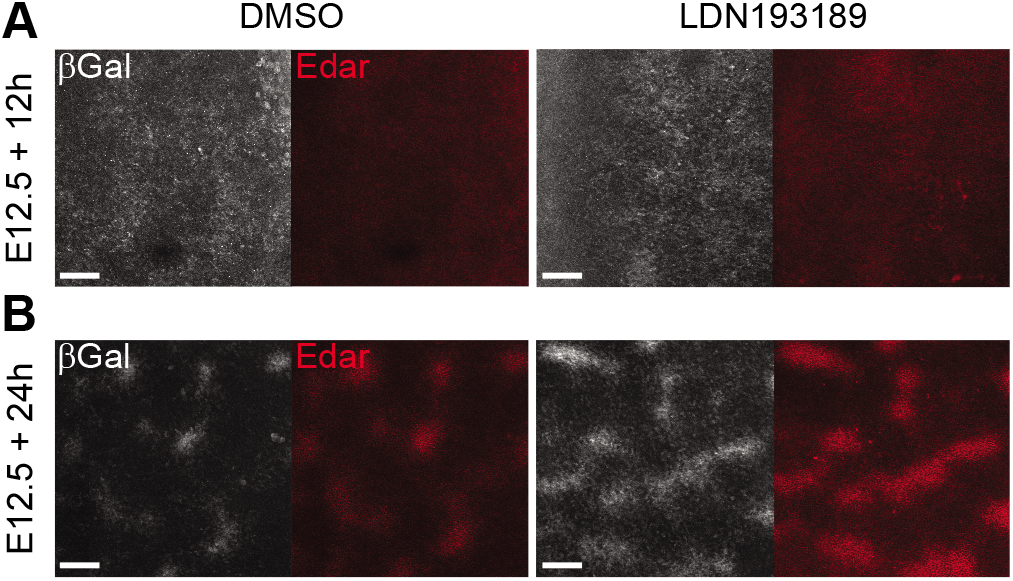
Inhibition of BMP signalling is insufficient to induce precocious hair placodes. (A, B) Maximum intensity projections of confocal image stacks of DMSO control and LDN193189 (1 µM) treated E12.5 Fgf20^βGal/+^ back skin explants after 12 hours (N = 0/10 for both) and 24 hours (N= 9/11 for both) of culture. Samples were stained with βGal and Edar antibodies to visualize nascent placodes. N indicates samples with induced placodes / total number of samples. Scale = 100 µm.

### Rspo1 signalling alone is insufficient to induce hair follicle induction

Given the essential role of epithelial Wnt/β-cat signalling in hair follicle induction (Zhang et al., 2009; Huelsken et al., 2001; Närhi et al., 2008; Zhang et al., 2008), we next focused our attention to potential activators of this pathway. As dermal deletion of *Wls*, a gene indispensable for Wnt ligand secretion, does not impair hair follicle induction (Chen et al., 2012), we hypothesized that the inductive signal could be another type of soluble Wnt agonist. R-spondins-1-4 (Rspo1-4) act synergistically with Wnt ligands to enhance Wnt/β-catenin signalling activity via Leucine Rich Repeat Containing G Protein-coupled Receptors 4-6 (Lgr4-6) (Nusse & Clevers, 2017; Steinhart & Angers, 2018). Intriguingly, *Lgr4* and *Lgr6* are expressed in the nascent primary hair placodes (Sennett et al., 2015; Tomann et al., 2016; Snippert et al., 2010) and loss of *Lgr4* leads to near total absence of primary hair placodes and greatly reduced total number of hair follicles in newborn mice (Mohri et al., 2008) implying an important role for Rspos in hair follicle induction.

Previous RNA-seq profiling of E14.5 back skin has indicated that *Rspo1* and *Rspo3*, but not *Rspo2* and *Rspo4*, are expressed in the mesenchyme (Sennett et al., 2015; Mok et al., 2019; Snippert et al., 2010). We first analysed the expression pattern of *Rspo1* and *Rspo3* in the back skin prior to hair follicle induction (E12.5), during hair follicle induction (E13.5), and at placode stage (E14.5) by RNA *in situ* hybridization. We did not observe focal expression of *Rspo1* at any stage, instead the signal appeared homogenous in the entire mesenchyme (Fig. 6A). Expression of *Rspo3* was unpatterned at E12.5 and E13.5, but at E14.5, a punctate expression in the DCs was evident, confirmed by wholemount *in situ* hybridisation (Fig. S4A), in line with RNA-seq profiling of DCs (Sennett et al., 2015). To quantify the expression levels of *Rspo1* and *Rspo3*, we performed qRT-PCR on E12.5-E14.5 mesenchymes. Interestingly, *Rspo1* expression was significantly upregulated between E12.5 andE13.5 (Fig. 6B), whereas the expression of *Rspo3* did not significantly alter between the stages (Fig. 6B). These data suggest that Rspo1 might be associated with hair follicle induction.

**Fig. 6.**
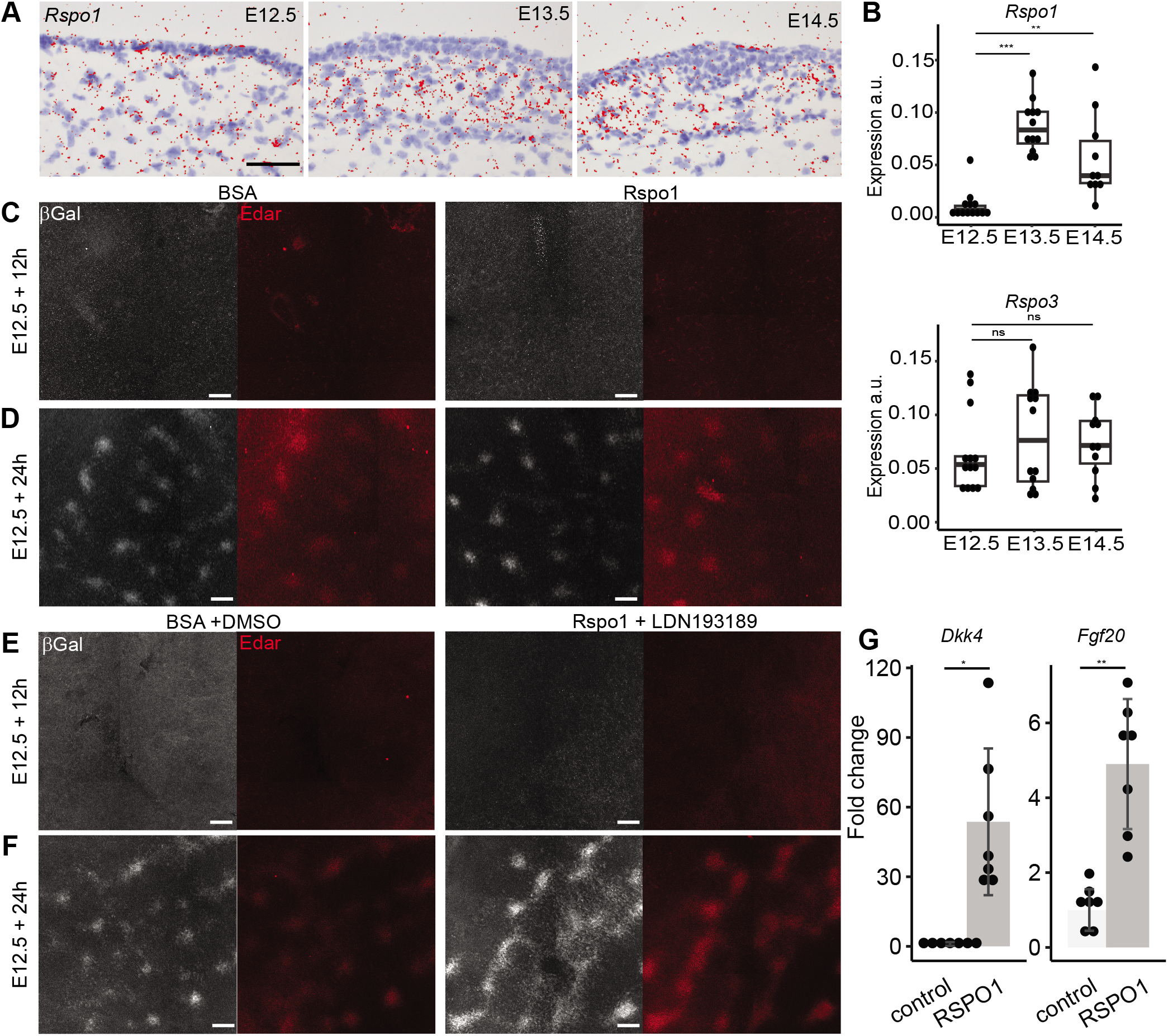
*Rspo1* is upregulated in dorsal mesenchyme upon hair follicle induction but is not sufficient to induce hair follicle development. (A) *Rspo1* expression in back skin of E12.5, E13.5, and E14.5 wild type embryos, detected by RNA *in situ* hybridization. Scale = 50 µm. (B) qRT-PCR quantification of *Rspo1* and *Rspo3* expression in back skin mesenchyme at E12.5, E13.5, and E14.5 normalized to *Gapdh* expression (Line indicates mean, whiskers variance). * P<0.05, **P<0.01, ***P<0.001, statistical significance was assessed with Student’s T-test. (C, D) Maximum intensity projections of confocal image stacks of BSA and Rspo1 (100 ng/ml) treated Fgf20^βGal/+^ E12.5 back skin explants after 12 hours (N = 0/9 for both) and 24 hours (N_DMSO_ = 13/14 and N_LDN_ = 14/15) of culture with βGal and Edar immunolabelling. Numbers indicate samples with induced placodes / total number of samples. Scale = 100 µm. (F, G) Maximum intensity projections of confocal image stacks of BSA + DMSO and Rspo1 (100 ng/ml) + LDN193189 (1µM) treated Fgf20^βGal/+^ E12.5 back skin explants after 12 hours (N = 0/7 for both) and 24 hours (N = 7/7 for both) of culture with βGal and Edar immunolabelling. (E) Fold change of expression of *Dkk4* (53.66 ± 29.27), and *Fgf20* (4.90 ± 1.61) in E12.5 back skin after 4 hours of 100 ng/ml Rspo1 treatment relative to vehicle control, measured by qRT-PCR (N=7, Mean ±SD). *P<0.05, **P<0.01. Statistical significance was assessed with Student’s T-test and Wilcoxon matched-pairs signed rank test for normally (*Fgf20*) and non-normally (*Dkk4*) distributed data, respectively.

In adult mice, exogenous recombinant Rspo1 injection leads to precocious activation of hair follicle stem cells during the resting phase of the hair cycle (Li et al., 2016). To examine whether Rspo1 is sufficient to induce hair placode formation in the embryonic skin, we cultured E12.5 back skin explants for 12 hours and 24 hours with 100 ng/ml Rspo1. At 12 hours, hair placodes had not formed in control or Rspo1-treated skins (Fig. 6C), whereas after 24 hours of culture, focal expression of placode markers was evident in both samples, indicative of placode induction (Fig. 6D). A higher Rspo1 concentration (500 ng/ml) did not induce precocious placodes either (Fig. S4B).

We also tested whether simultaneous activation of Wnt/β-catenin signalling via supplementation of Rspo1 and inhibition of BMP signalling together would be sufficient to advance hair placode formation. To this end, we cultured E12.5 back skin for 12 hours in media supplemented with 100 ng/ml Rspo1 and 1 µM LDN193189, or BSA and DMSO as vehicle controls, but observed no formation of hair placodes in either condition (Fig. 6G). When hair placodes emerged after 24 hours of culture, they were more irregular and even stripe-like in samples supplemented with Rspo1 and LDN193189 (Fig. 6F), similar to those treated with LDN193189 alone (Fig. 5B). Taken together, *ex vivo* manipulation of the developing skin with Rspo1 and LDN193189, alone or together, was not sufficient to induce precocious hair placode formation.

These results raised the question whether exogenous Rspo1 can induce a Wnt signalling response in the naïve epidermis, prior to placode formation. To assess this, we treated E12.5 embryonic skin with 100 ng/ml recombinant Rspo1 in hanging drops for 4 hours and measured the signalling response by analysing expression of placode markers and Wnt/β-catenin target genes *Fgf20* and *Dkk4* (Bazzi et al., 2007; Fliniaux et al., 2008; Huh et al., 2013) by qRT-PCR. We observed a significant increase in the expression of both *Dkk4* (53.66-fold ± 29.27) and *Fgf20* (4.90-fold ± 1.61) upon Rspo1 treatment (Fig. 6E) indicative of a positive Wnt signalling response.

## Discussion

To date, the source of the hair follicle inductive signal has remained unknown. Here we show that the hair-forming back skin mesenchyme induces hair development in the normally hairless plantar epithelium at developmental stages after DC formation (E14.5), at the time of epithelial thickening (E13.5), as well as prior to DC formation (E12.5). Even the E12.5 *Fgf20* null mesenchyme that is devoid of primary and secondary hair follicle DCs induced hair follicle formation in the normally glabrous epithelium. These data confirm that the mesenchyme is the source of the inductive cue(s) even prior to any molecular or morphological signs of hair follicle induction. This contrasts with the observations of the developing tooth where the tooth inductive capacity resides initially in the epithelium, but shifts to the mesenchyme concomitant with the condensation of the dental mesenchyme (Mina & Kollar, 1987; Lumsden, 1988). In our recombination experiments, the back skin epithelium failed to induce hair follicle formation at all stages analysed.

In developing chicken embryos, a global increase in mesenchymal cell density precedes feather follicle induction (Wessells, 1965; Ho et al., 2019). We found no evidence for a similar change in the upper dermis of mouse embryos at the time of hair placode formation suggesting that alterations in cellular gene expression levels rather than cell density govern hair follicle induction. The identity of the hair follicle inductive signal remains elusive to date. However, all available evidence points to positive and negative roles for the Wnt/β-catenin and Bmp signalling pathways, respectively (Biggs & Mikkola, 2014). Here, we assessed the ability of the Bmp receptor inhibitor LDN193189 and Wnt agonist Rspo1 to induce hair follicle development. Although suppression of Bmp signalling changed the regular arrangement of the primary hair placodes into a stripe-like pattern, in line with its proposed involvement in hair follicle patterning (Mou et al., 2006; Glover et al., 2017), it did not result in precocious placode formation. *In vivo,* hair follicle formation is suppressed when *Noggin* is ablated (Botchkarev et al., 1999, 2002), while its overexpression leads to formation of excess and ectopic follicles (Plikus et al., 2004). In cultured skin explants, exogenous Bmp4 prevents placode formation (Mou et al., 2006; Pummila et al., 2007). Together these results show that Bmp signalling plays a central role in hair follicle induction and patterning, yet its inhibition alone appears to be insufficient to trigger hair follicle development. In developing skin, downregulation of epithelial Bmp activity might be achieved via concomitant downregulation of pathway agonists (such as Bmp4) and upregulation of soluble antagonists, many of which are expressed in the mesenchyme at the time of hair follicle induction (Jacob et al., 2022 preprint; Sennett et al., 2015).

Augmentation of Wnt signalling via application of Rspo1 alone or together with LDN193189 did not induce precocious hair follicle development either, although we did observe a significant upregulation of hair placode marker genes *Dkk4* and *Fgf20* upon Rspo1 treatment at E12.5, providing evidence that the naïve epidermis is responsive to Rspo1 prior to hair follicle induction. While the failure in hair placode induction may indicate that additional signalling pathways are necessary, alternative explanations are also plausible. The periodic organization of hair follicles is thought to arise via the Turing reaction-diffusion system (Andl et al., 2002; Sick et al., 2006; Mou et al., 2006; Glover et al., 2017). This self-organizing model posits that spatial patterns result from the interaction of diffusible signals that minimally consist of a pair of interacting diffusible chemical signals: a short-range, self-enhancing activator and an activator-induced, longer-range inhibitor (Turing, 1952; Gierer & Meinhardt, 1972; Kondo, Miura & Turing, 2010). As this system leads to focal production of both the activator and the inhibitor, supplying the activator (such as Rspo1) in the culture medium might initially flatten the Wnt activity pattern and thereby prevent follicle induction. It is also possible that Rspo1 alone is not sufficient to increase Wnt signalling activity above a critical threshold. We speculate that to achieve this threshold, combined upregulation of mesenchymally produced agonists (such as Rspo1) and downregulation of antagonists (such as Dkks and Secreted Frizzled-related proteins, Sfrps) might be necessary. Indeed, a recent transcriptomic profiling study revealed strong downregulation of *Dkk2* in the upper dermis at the time of hair follicle induction (Jacob et al., 2022 preprint). *Dkk2* is also expressed at higher levels in the non-hairy regions of the embryonic skin compared to the hair-forming regions, and its deletion is sufficient to induce ectopic hair follicle formation in the plantar skin (Song et al., 2018) and cornea (Mukhopadhyay et al., 2006), underscoring the significance of diffusible mesenchymal Wnt inhibitors in hair follicle induction.

Taken together, our study complements the tissue recombination studies of inductive signalling in hair follicle development by establishing that the primary inductive cue arises from the mesenchyme. While the identity of the signal remains elusive, our results from back skin mesenchyme-oral epithelium recombination experiments reveal that the hair inductive cue is readily interpreted also by a tissue of non-skin origin resulting in the formation of functional hair follicles. Recombination of DC-containing whisker pad mesenchyme with the dental epithelium has demonstrated that under the inductive signals of the DC, the dental epithelium switches from the tooth-forming program to one generating whiskers (Kollar, 1970, 1966). As the oral epithelium used in our experiments was devoid of tooth placodes, our interpretation is that E12.5 back skin mesenchyme induced hair follicles *de novo*, rather than by transforming an existing tooth primordium into a hair-producing organ. Instead, genetic activation of epithelial Wnt signalling induces ectopic epithelial invaginations and teeth, not hair follicles, in the oral epithelium (Närhi et al., 2008; Zhang et al., 2008; Wang et al., 2009). This together with our results imply that hair follicle identity is governed by mesenchymal cues other than those merely intensifying epithelial Wnt activity and placode formation. Importantly, our results demonstrate that hair follicle inductive capacity is an intrinsic property of the naïve skin mesenchyme even prior to formation of the dermal condensate.

## Materials and Methods

### Mouse lines

| Mouse line | Source or reference | Identifiers |
| --- | --- | --- |
| wildtype NMRI | Laboratory animals centre,<br>Helsinki institute of Life<br>Science, University of<br>Helsinki |  |
| Fgf20<sup>β-Gal/β-Gal</sup> | PMID: 23431057 | RRID:MGI:5425887 |
| K17-GFP | PMID: 16055733,<br>Jackson Laboratory | Stock: 023965,<br>RRID:IMSR_JAX:023965 |
| R26R<sup>tdTomato</sup> | Jackson Laboratory | Stock 007914 |
| Pgk-Cre | PMID: 9608738,<br>Jackson Laboratory | RRID:IMSR_JAX:020811 |
| R26R<sup>tdTomato</sup> | PMID:20023653,<br>Jackson Laboratory | Stock 007914,<br>RRID:IMSR_JAX:007914 |
| R26R<sup>mTmG</sup> | Jackson Laboratory | Stock 007576,<br>RRID:IMSR_JAX:007576 |
| TCF/Lef1:H2B-GFP | Jackson Laboratory | Stock 013752,<br>RRID:IMSR_JAX:013752 |

### Antibodies

| Antibody | Producer | Identifiers |
| --- | --- | --- |
| goat, Sox2 | Santa Cruz | SC-17320 |
| rabbit, Sox2 | Stemgent | 09-0024 |
| goat, Lhx2 | Santa Cruz | SC-19344 |
| rat, EpCAM | BD Pharmingen | 552370 |

### Reagents and kits

| Reagent | Producer | Identifiers |
| --- | --- | --- |
| RSPO1, Recombinant Mouse R-spondin1 (CHO-exp) Protein, CF | R&D Systems | cat. 7150-RS-050/CF |
| Stemolecule™ LDN193189 | Stemgent (Lexington, MA) | cat. 04-0074 |
| RNEase Plus Mini Kit | Qiagen, GmbH | REF. 74126 |
| iScript gDNA Clear cDNA Synthesis Kit | Bio-Rad (Hercules, CA) | cat. 1725035 |
| Dkk4, Mmu, FAM Probe Assay | Bio-Rad (Hercules, CA) | cat. 10031225 |
| GAPDH, Mmu, Cy5 Probe Assay | Bio-Rad (Hercules, CA) | cat. 10031231 |
| Fgf20, Mmu, TEX615 Probe Assay | Bio-Rad (Hercules, CA) | cat. 10031234 |
| iTaq Universal Probes Supermix | Bio-Rad (Hercules, CA) | cat. 1725131 |
| Höchst33342 | Invitrogen | REF. H3570 |
| Proteinase K | Roche | REF 3115879001 |

### Ethics statement

All mouse studies were approved and carried out in accordance with the guidelines of the Finnish national animal experimentation board.

### Mouse lines

R26R^mTmG^ mouse line was obtained from Jackson Laboratory and, as previously described in (Lan et al., 2023 preprint), crossed to Pgk-Cre to obtain mice where R26R^mTmG^ is recombined to R26R^mG^, and constitutively and heritably expressed from the Rosa26 locus in every cell. R26R^tdTomato^ line was obtained from Jackson Laboratory and crossed to Pgk-Cre (also from Jackson Laboratory, Stock: 020811) to obtain mice where tdTomato is constitutively and heritably expressed from the Rosa26 locus in every cell. In hanging drop experiments, wild type NMRI embryos were used.

### Tissue recombination

The back skins were dissected from embryos of the indicated stages and genotypes. To obtain the plantar skin, the entire distal part of the limb was dissected. To obtain the diastema tissue from lower jaws, the mandible was dissected and trimmed to contain only the lingual surface of the mandible with the incisor and molar tooth primordia to mark borders of the diastema. The dissected tissues were treated with 2.5 U/ml Dispase II (Roche) in PBS, +4°C for the following times: E12.5 dorsal and plantar skin 20 min, E13.5 back skin 25-30 min, E14.5 40 min, oral tissue 8 min. The tissues were then allowed to rest in culture medium at room temperature for 30 min. The plantar skin was dissected from the rest of the limb, and all epithelial and mesenchymal tissues separated using forceps and needles. Epithelial and mesenchymal tissues were recombined so that the epithelium was placed on top of the mesenhcyme. Tissue separation efficiency was assessed by microscopy. The separation was always efficient, yet it was not possible to avoid the transfer of single mesenchymal cells with the epithelium.

### Ex vivo tissue culture

Tissues were cultured in the air-liquid interface in a Trowell-type setup as described earlier (Närhi & Thesleff, 2010) using DMEM, 10% FBS, 1% penicillin-streptomycin (PS), both supplemented with 10 U/ml penicillin, 10 µg/ml streptomycin, except with recombinations including oral tissues when DMEM:F12 (1:1) supplemented with 10% FBS, 1% PS, and 100 ng/ml ascorbic acid was used. Media were exchanged every other day. When indicated, the media were further supplemented with RSPO1 (R&D Systems), LDN193189 (Stemgent, Lexington, MA) at concentrations indicated in the text, and BSA and DMSO used as vehicle. At the end of the culture, tissues were fixed in 4% paraformaldehyde in PBS for immunofluorescence staining 1 h at room temperature, and washed with PBS.

### Vibratome sectioning

To make transversal vibratome sections of E12.5 and E13.5 embryos, the embryos were decapitated and fixed in 4% paraformaldehyde in PBS overnight at +4°C. The embryos were then embedded in 2% low-melting point agarose (Thermo Scientific, Vilnius, Lithuania) in PBS, sectioned to 250 – 300 µm sections using Microm HM650 V vibratome, and the sections were collected in PBS prior to immunostaining.

### Immunostaining and confocal microscopy

The fixed tissues were washed in PBS and stained with antibodies using standard protocols. In brief, samples were blocked and permeabilized using PBS, 10% normal donkey serum, 0.3% Triton-X, room temperature, 1 h, followed by overnight incubation at +4°C with the primary antibodies diluted in the blocking solution: Sox2 (1:250), LHX2 (1:400), EpCAM (1:500), β-Galactosidade (1:1000), and Edar (1:400). All samples we co-stained with 1:2000 Hoechst33342. The samples were further washed with several changes of PBS over 6 hours at room temperature prior to incubation with AlexaFluor conjugated secondary antibodies diluted 1:400 in PBS overnight at +4°C. The following day, the samples were washed with several changes of PBS over minimum of 4 h at room temperature and mounted on microscopy slides with Anti-Fade Fluorescense Mounting Medium (Abcam).

The samples were imaged using Zeiss LSM 700 laser scanning confocal microscope equipped with PMT using Plan-Apochromat 10x/0.45, LD LCI Plan-Apochromat 25x/0.8 Imm Corr, and LCI Plan-Neofluar 63x/1.30 Imm Corr objectives. To analyse dermal cell density, image stacks of the vibratome sections were obtained using the 63x objective at 0.5 µm intervals. To analyse treated skin explants for placode induction images were obtained with a 10x objective at 5-10 µm intervals.

### Image analysis

To quantify mesenchymal cell densities, confocal image stacks of vibratome sections were processed and analysed using Imaris 10.0 and R softwares. Firstly, the attenuation of signal in the sample was determined by measuring average Hoechst33342 signal intensities on the top and bottom z-sections of the stack in 5 - 10 measuring points made on the nuclei. The attenuation correction function of Imaris was then used on the channels of the image. The epithelial surface was created manually based on the anti-EpCAM staining and a distance transformation was made to delimit the 80 µm of the mesenchyme to epithelium. The nuclear Hoechst33342 signal within this volume was then used to segment the nuclei using the machine learning Labkit plugin (Arzt et al., 2022) by classifying Hoechst33342 and background pixels. Distance of the cell to epithelium was determined as the intensity of the distance transformation channel at the centre and the TCF/Lef1:H2B-GFP intensity as the mean intensity of the GFP channel of each segmented nucleus. The volumes at each 1 μm distance interval were determined using the auxiliary ImageJ histogram function, by quantifying the number of pixels with respective values of the distance transformation channels within the delimited 80 μm from the epithelium volume. The densities were calculated, and statistical testing was performed using R-studio software.

For *ex vivo* cultured samples, the confocal image stacks were visualised as maximal intensity projections and placode induction was determined as patterned upregulation of Fgf20^βGal^ reporter and Edar expression.

### Hanging drop culture, RNA extraction, and qRT-PCR

The short-term treatment of tissue samples in the hanging drop culture was performed as previously described (Biggs et al., 2018). Briefly, in two independent experiments E12.5 skins were dissected from NMRI embryos and cut in half along the dorsal midline. The explants were then placed individually in 40 μl droplets of DMEM, 10% FBS, 1% penicillin-streptomycin supplemented with either 100 ng/ml RSPO1 (R&D Systems) or the respective volume of 0.1% BSA in PBS for vehicle control under a culture dish lid that was turned on a culture dish, and maintained for 4 hours at 37°C, 5% CO_2_. RNA was extracted from the samples using the RNEasy Plus Mini Kit (Qiagen GmbH, Hilden, Germany) according to manufacturer’s instructions. cDNA was synthesized from 1 µg RNA using iScript gDNA Clear cDNA Synthesis Kit (BIORAD, Hercules, CA), according to manufacturer’s instructions and the cDNA was diluted to 5 ng/µl.

For analysis of *Rspo1* and *Rspo3* gene expression, the back skin was dissected from wild type NMRI embryos at E12.5 (N=13), E13.5 (N=12), and E14.5 (N=10), treated with Dispase II as described above to separate the mesenchyme from the epithelium. The mesenchyme was collected, and the RNA extracted, and cDNA translated as above.

For embryonic skin samples treated with Rspo1, qRT-PCR was performed using gene-specific PrimePCR Probe Assay (Bio-Rad, Hercules, CA) for mouse *Dkk4*, *Fgf20* and *Gapdh* multiplexed in 20 µl reactions according to manufacturer’s instructions with 25 ng of template. Reactions were performed in triplicate wells with the CFX96 Real-Time System (Bio-Rad, Hercules, CA) with the protocol described earlier (Biggs et al., 2018). The fold changes were calculated with the Δ ΔCT method (Livak, Schmittgen & Methods, 2001), by normalizing to average of reference gene *Gapdh* in all controls. The normal distribution of statistical difference was analysed using paired Student’s T-test.

Expression of *Rspo1* and *Rspo3* in the embryonic skin mesenchyme was analysed with primer-based RT-qPCR using Fast-SYBR Green Master Mix (Thermo Fisher Scientific, Vilnius, Lithuania) in 20 μl reactions with 15 ng template and 0.5 µM of each primer. *Rspo1* primers were designed using template NM_138683.2: forward AGAGACAGAGGCGGATCAGTG and reverse CAGAATGAAGAGCTTGGGCG, and *Rspo3* using template NM_028351.3: forward GTCAGTATTGTACACTGTGAGGC and reverse CCGTGTTTCAGTCCCCCTTT. *Gapdh* primers were: forward CTCGTCCCGTAGACAAAATGG and reverse AGATGGTGATGGGCTTCCC. Reactions were run in triplicate as above. The relative expression was calculated using the ΔCT method by normalizing to *Gapdh* and analysed with Student’s T-test.

### In situ hybridization

For whole-mount RNA *in situ* hybridisation, E14.5 wild type NMRI embryos were decapitated and fixed in 4% PFA in PBS overnight at 4°C, and then dehydrated in a series of methanol. The hybridization was performed manually using a protocol described before (Biggs *et al*. 2018). Antisense RNA probes were designed to target *Rspo1* (template NM-138683.2, base pairs 924-1359) and *Rspo3* (NM-028351.3, base pairs 975-1448). The samples were imaged using Zeiss AxioZoom.V16 stereomicroscope with PlanZ 1.0x/C objective and Axiocam 305 color camera.

For radioactive *in situ* hybridization on paraffin sections, the E12.5 (N=4), E13.5 (N=5), and E14.5 (N=5) embryos were decapitated, fixed in 4% PFA overnight at 4°C, processed into paraffin blocks using standard protocols, and section in 5 µm sagittal sections. The *in situ* hybridization was performed according to standard protocols (Huh et al., 2013). Imaging and image processing was performed as previously described (Biggs et al., 2018).

## Acknowledgements

We thank Ms. Raija Savolainen, Ms. Riikka Santalahti, and Ms. Merja Mäkinen for excellent technical assistance, Ms. Verdiana Papagno for assistance with *in situ* hybridization, Dr. Qiang Lan for crosses of mice, Dr. Kimmo Tanhuanpää for expert advice on image analysis, and past and present members of the Mikkola, Jukka Jernvall, and Anamaria Balic labs for stimulating discussions. We thank Dr. Leah Biggs for critical reading of the manuscript. Mouse studies were carried out with the support of HiLIFE Laboratory Animal Center Core Facility, University of Helsinki. Confocal microscopy was conducted at the Light Microscopy Unit, Institute of Biotechnology, supported by HiLIFE and Biocenter Finland.

## Author Contributions Statement

Otto Mäkelä: Conceptualization, Methodology, Investigation, Formal Analysis, Writing – Original Draft, Writing – Review & Editing, Visualization

Marja Mikkola: Conceptualization, Methodology, Writing – Review & Editing, Supervision, Project administration, Funding acquisition

---

**Fig S1 (Related to Fig. 2).**
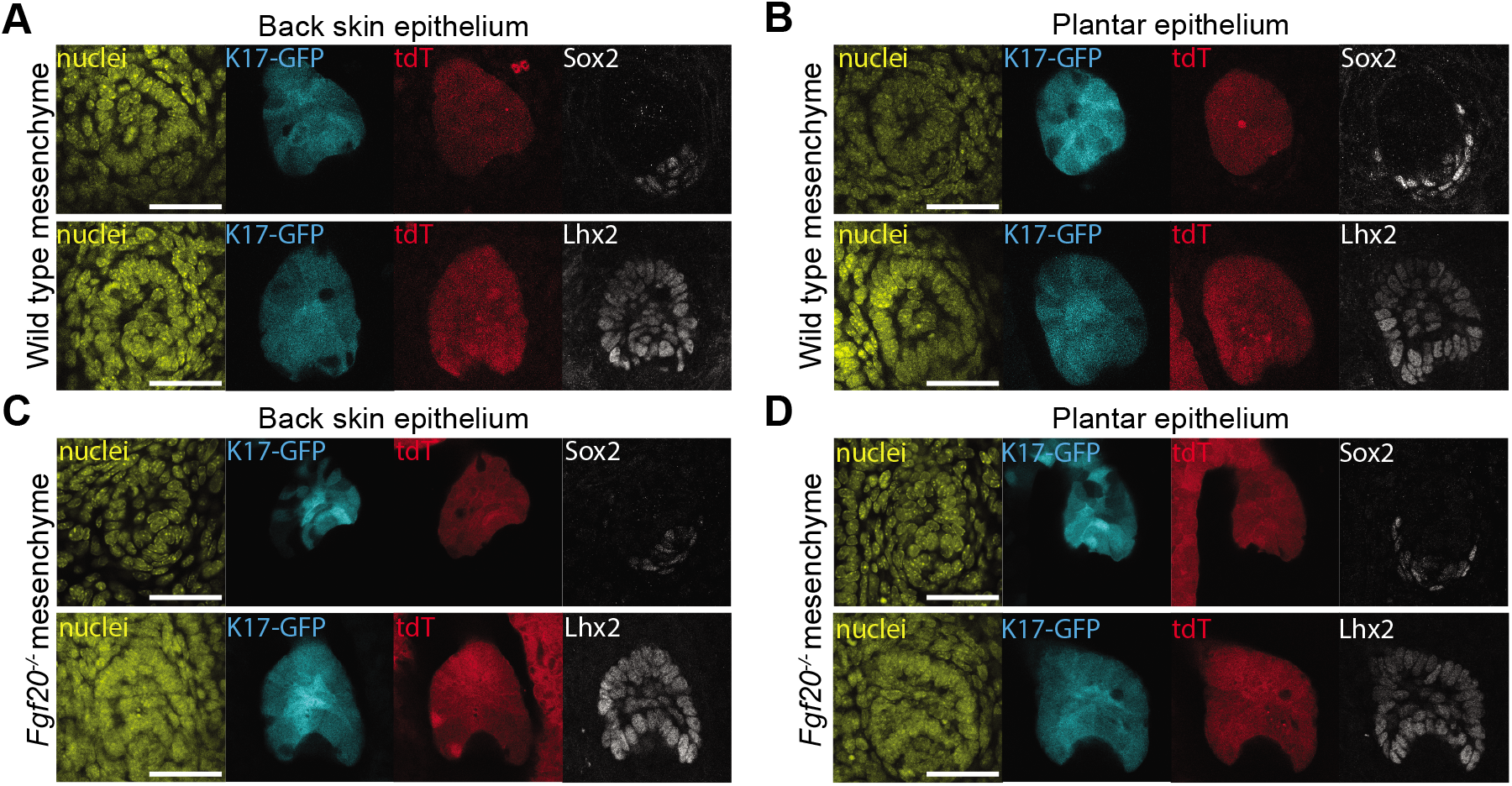
Appendages induced in recombinants of E12.5 wild type or Fgf20_-/-_ back skin mesenchyme and E12.5 plantar epithelium express hair follicle markers. (A, B) Single channels of the confocal optical sections of recombinants of E12.5 wild type mesenchyme and back skin control (A) and plantar epithelium (B) from E12.5 R26R^floxedTOM^;K17-GFP embryos stained with Sox2 or Lhx2. (C, D) Single channels of the confocal optical sections of recombinants of E12.5 Fgf20^-/-^ mesenchyme and back skin (C) and plantar epithelium (D) from E12.5 R26R^floxedTOM^;K17-GFP embryos stained with Sox2 or Lhx2. Nuclei were stained with Hoecsht33342. Scale 20 µm.

**Fig S2 (Related to Fig. 3).**
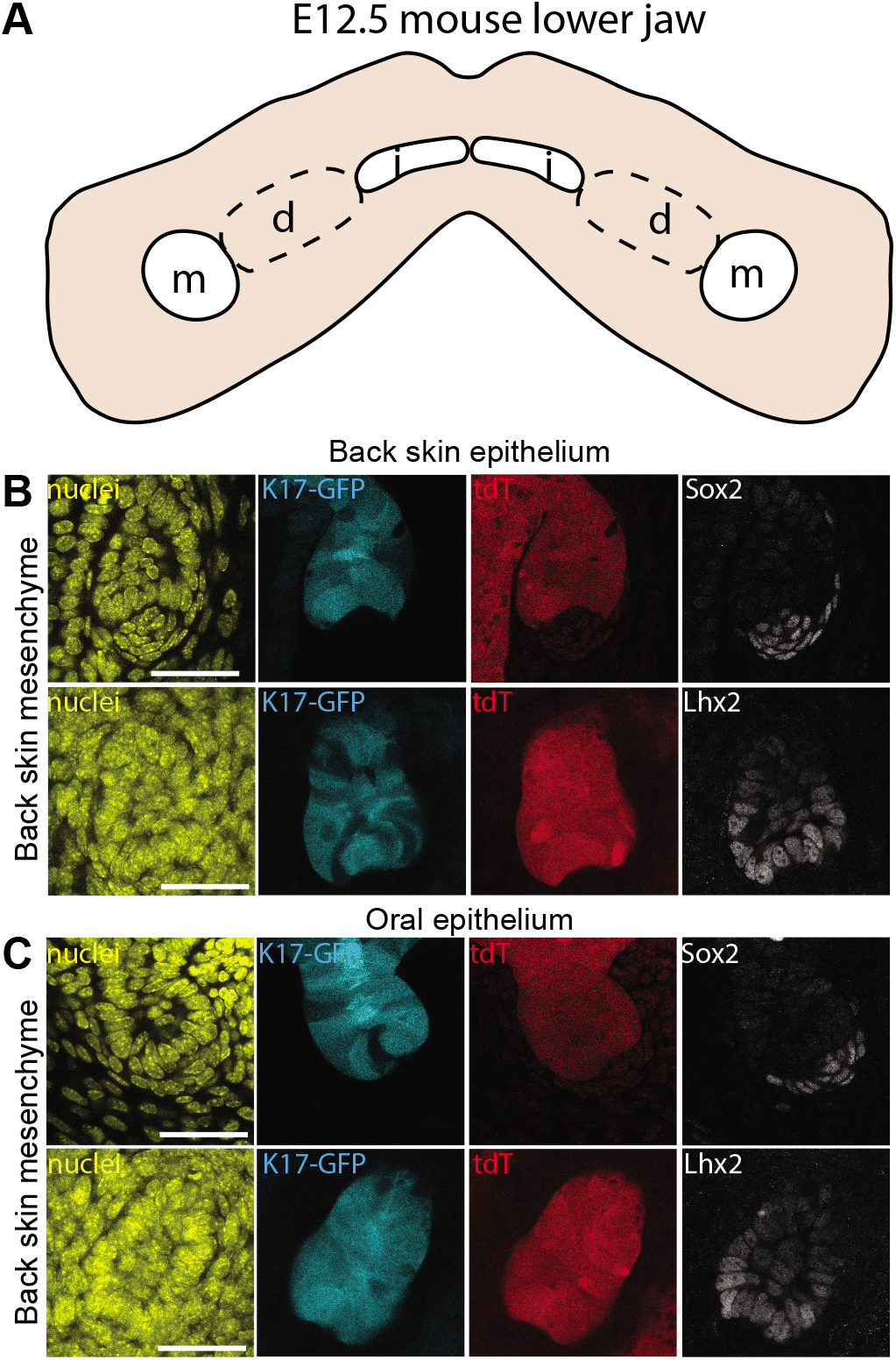
Appendages induced in recombinants between E12.5 back skin mesenchyme and oral epithelium express hair follicle markers. (A) A schematic view of the E12.5 mouse lower jaw. White areas indicate incisor (i) and molar (m) primordia and area marked by the dashed line indicates diastema (d) region used in the tissue recombination experiments. (B, C) Single channels of the confocal optical sections from the appendages of the homotypic (B) and heterotypic (C) recombinations in Fig 3. Scale 20 µm.

**Fig S3 (Related to Fig. 4).**
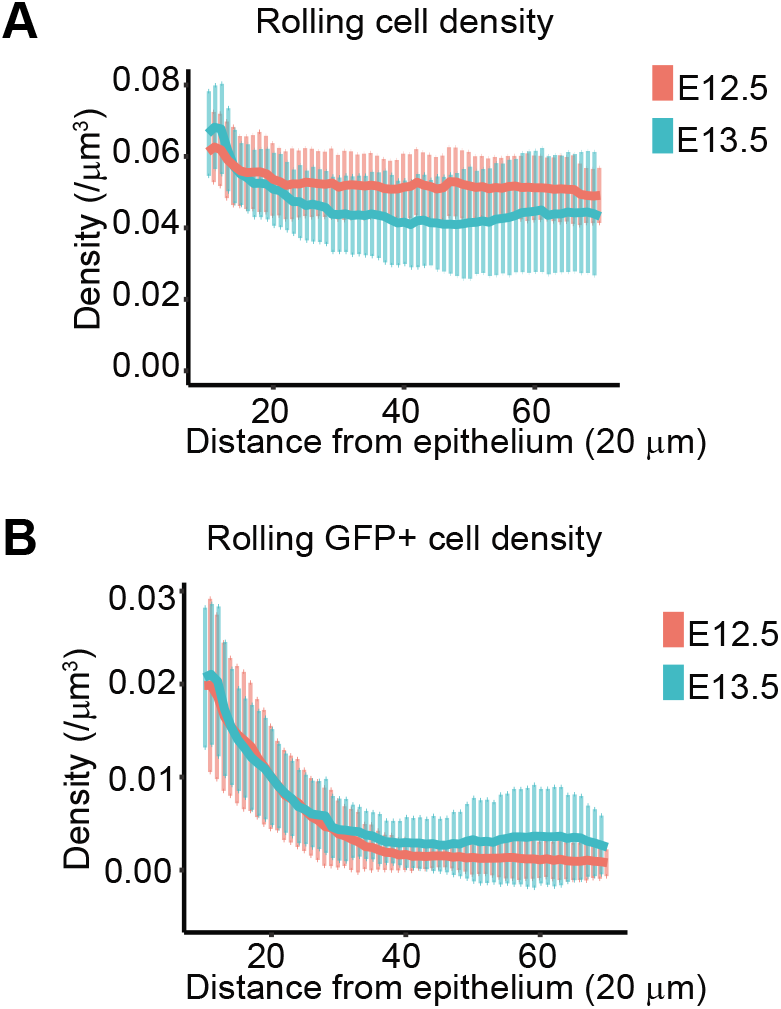
Mesenchymal cell density does not change during hair follicle induction. (A) Mean rolling densities of all mesenchymal cells within 20 µm windows at 1 µm intervals (nuclei/µm^3^, Mean ±SD) at E12.5 (red) and E13.5 (blue). (B) Mean rolling densities of TCF/Lef1:H2B-eGFP+ cells within 20 µm windows at 1 µm intervals (nuclei/µm^3^, Mean ±SD) at E12.5 (red) and E13.5 (blue).

**Fig S4 (Related to Fig. 6).**
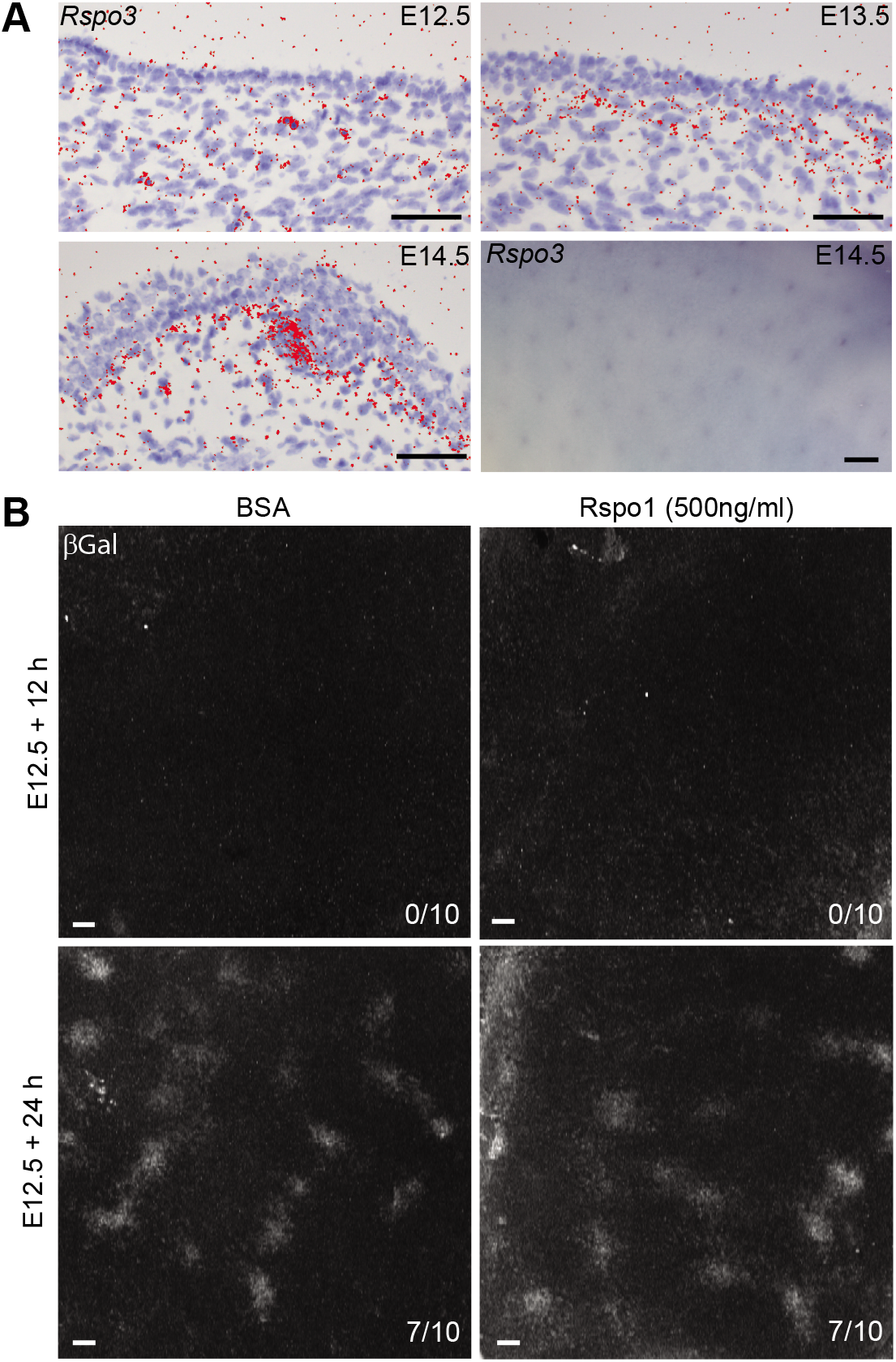
*Rspo3* is expressed in dermal condensates. (A) *Rspo3* expression in back skin of E12.5 (N=4), E13.5 (N=5), and E14.5 (N=5) wild type embryos, detected by RNA *in situ* hybridization. *Rspo3* whole mount *in situ* hybridization on E14.5 embryo (N=9). (B) Maximum intensity projection of confocal image stacks of BSA vehicle control and Rspo1 (500 ng/ml) treated Fgf20^βGal/+^ E12.5 back skin explants after 12 and 24 hours of culture with βGal immunolabelling. Numbers indicate samples with induced placodes / total number of samples. Scale = 50 µm, except wholemount ISH of *Rspo3* 200 µm.

